# Brain-machine interface learning is facilitated by specific patterning of distributed cortical feedback

**DOI:** 10.1101/2019.12.12.873794

**Authors:** A. Abbasi, H. Lassagne, L. Estebanez, D. Goueytes, D. E. Shulz, V. Ego-Stengel

**Author notes:** Equal contribution (A. Abbasi, H. Lassagne, L. Estebanez). Equal contribution (D. E. Shulz, V. Ego-Stengel).

## Abstract

Neuroprosthetics offer great hope for motor-impaired patients. One obstacle is that fine motor control requires near-instantaneous, rich somatosensory feedback. Such distributed feedback may be recreated in a brain-machine interface using distributed artificial stimulation across the cortical surface. Here, we hypothesized that neuronal stimulation must be contiguous in its spatiotemporal dynamics in order to be efficiently integrated by sensorimotor circuits. Using a closed-loop brain-machine interface, we trained head-fixed mice to control a virtual cursor by modulating the activity of motor cortex neurons. We provided artificial feedback in real time with distributed optogenetic stimulation patterns in the primary somatosensory cortex. Mice developed a specific motor strategy and succeeded to learn the task only when the optogenetic feedback pattern was spatially and temporally contiguous while it moved across the topography of the somatosensory cortex. These results reveal new properties of sensorimotor cortical integration and set new constraints on the design of neuroprosthetics.

**Teaser:** Targeting a brain-machine interface feedback to a cortical map reveals unexpected constraints.

## Introduction

Accurate limb control requires somatosensory feedback. For instance, local peripheral anesthesia blocking afferent tactile sensation in humans reduces dexterity and impairs fine motor control of the hand (*1, 2*). Similarly, cortical inactivation of somatosensory cortex in animals has profound effects on motor control (*3, 4*). The critical role of somatosensory feedback has also been described in studies of patients that suffer from severe tactile or proprioceptive deficits. These patients learn to rely extensively on visual feedback, but remain unable to manage normal motor control (*5–7*).

In the context of neuroprosthetics, proprioceptive and touch-like feedback originating from the prosthesis improves control (*8*), and enables texture-like percepts that cannot be obtained through visual feedback alone (*9*). Such artificial touch-like information has been provided through direct activation of the cerebral cortex via electrical stimulation (*8*, *10–14*) or optogenetics (*15*, *16*). Beyond the choice of the neuronal stimulation technology, an important challenge is the design of the geometry and dynamics of the feedback patterns used to provide relevant sensory feedback information.

The design of artificial sensory feedback is particularly critical for replicating the functionality of a spatially distributed sense such as touch (*17*). Temporal modulation of one single stimulation channel, such as realized by optogenetic stimulation of S1 in the BMI experiments of Prsa and collaborators (2017), cannot suffice in this case. Rather, many independent channels of stimulation will be necessary to convey tactile information arising from different peripheral locations. Indeed, recent approaches have implemented simultaneous artificial stimulations at multiple locations in the somatosensory cortex (*18*, *19*). However, it remains unclear if any arbitrary feedback pattern can be applied, or if the somatosensory-motor cortical areas can only integrate efficiently inputs with a specific type of spatio-temporal structure matched to the classical somatosensory topography (*20*).

Here, we take advantage of the well-known whisker system of the mouse to explore this question (*21*, *22*). Anatomically, the primary somatosensory cortex (S1) of the mouse snout is organized into distinct columns, called barrels, that each receive dominant inputs from one corresponding whisker. These inputs combine with dense subcortical and intracortical lateral connectivity (*23*, *24*) and give rise to rich encoding of complex multiwhisker features, which can be found at the level of individual neurons as well as in the cortical map (summarized in (*25*). Specifically, given the strong tuning of S1 neurons to the direction of bar-like multiwhisker deflections on the snout (*26*, *27*), and their tuning to progressive movement of objects across the whiskerpad (*28*), we hypothesized that stimulations that generate spiking activity in spatio-temporally contiguous barrel cortex locations may be more efficiently integrated into motor control, compared to spatio-temporally scattered patterns of cortical activation. This hypothesis is further supported by the fact that mice generate sequences of consecutive whisker deflections in the rostro-caudal direction during the exploration of their environment (*29*), suggesting that such stimuli are behaviorally relevant.

We tested this hypothesis by training mice to control a virtual cursor using the modulation of the activity of a few neurons, called Master neurons, recorded in the whisker area of the primary motor cortex (M1) (*30*). Mice received online one of 5 different spatiotemporal patterns of cortical feedback generating spiking activity in S1. These patterns ranged from a sweeping, bar-like feedback where the barrels that were simultaneously or sequentially activated were all contiguous, up to a spatiotemporally fully randomized pattern, also including a condition without feedback. The inputs were delivered on the surface of the cortex by photostimulation of subsets of always 5 barrels among the 22 most caudal barrels. Importantly, we focused on the impact of changes in the structure of patterned stimulation, while the total surface area, intensity and temporal frequency of stimulation remained always constant. We found that learning was largely dependent on the structure of the feedback and was highest in the bar-like feedback condition, where the photostimulated barrels are spatially and temporally contiguous. Learning in this specific condition revealed active motor control, as the neuronal activity that drove the virtual cursor reorganized and became dominated by one of the Master neurons.

## Results

### Patterned optogenetic feedback on S1 enables learning in a brain-machine interface

We implanted a total of 16 mice with a chronic, closed-loop brain-machine interface consisting of a head fixation bar, chronic silicon tetrodes in layer 5 of whisker M1 (Fig. 1A,B, Fig. S1, S2) and a chronic optical window over the S1 area (see Methods).

**Fig. 1.**
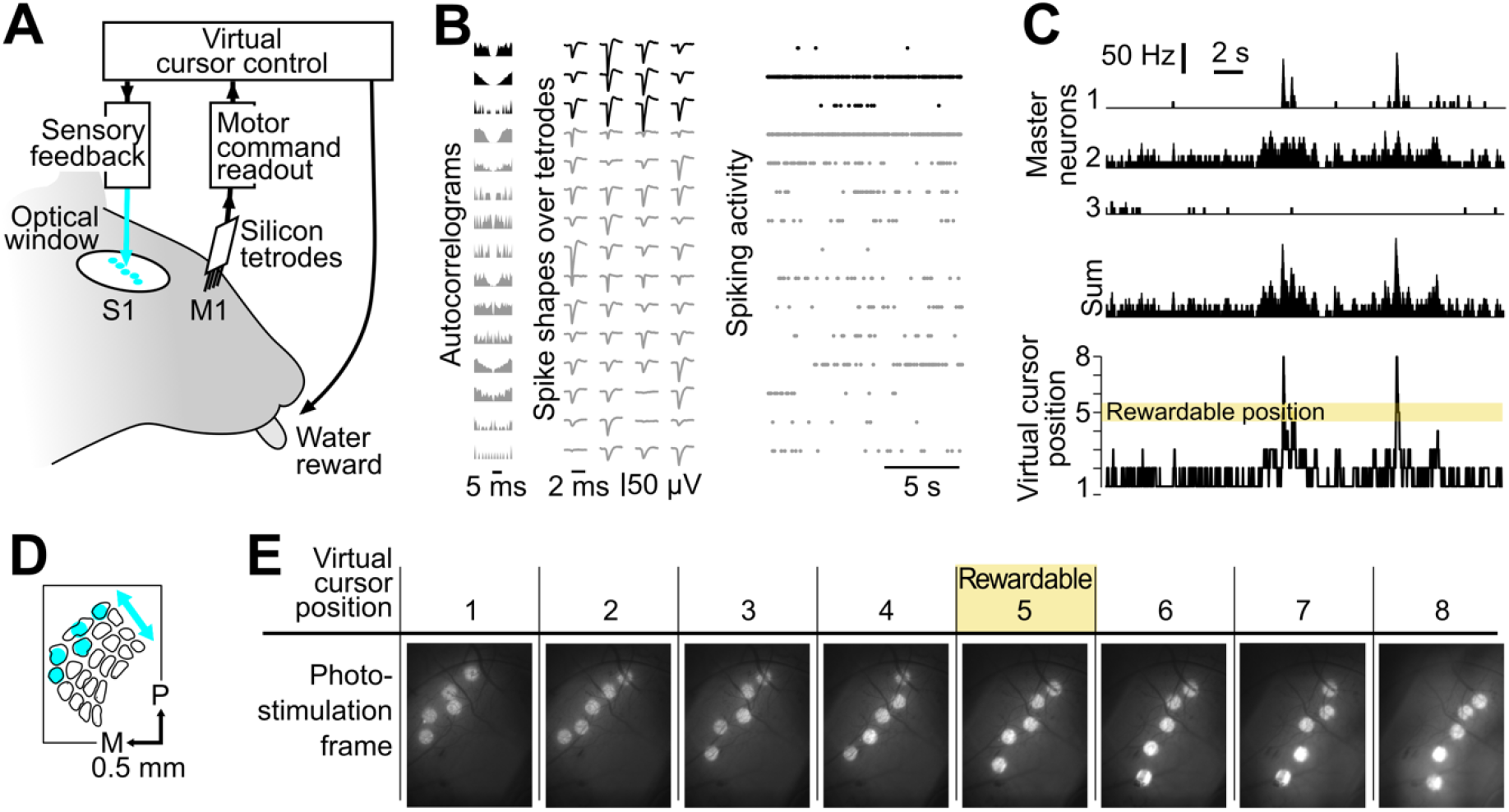
Mice controlled a virtual cursor using whisker M1 neuronal activity while online optogenetic feedback was delivered to whisker S1. **(A)** General view of the closed-loop interface. The mice were head-fixed. A chronic silicon probe in M1 read out spiking activity and a chronic optical window over S1 allowed delivery of a photostimulation feedback. **(B)** Action potentials from 15 single units obtained during baseline activity in M1. The autocorrelograms (left), the spike shapes on the tetrodes (middle) and the spiking activity in time (right) are shown for each single unit. Black: Master neurons that are selected to control the virtual cursor. Gray: Neighboring neurons recorded simultaneously. **(C)** Example Master neurons activity and corresponding virtual cursor position. Top: Time histograms of the 3 Master neurons activity. Middle: Sum of their activity. Bottom: Position of the virtual cursor computed from the summed activity of the Master neurons. The virtual cursor must be in position 5 for the mouse to obtain a reward by licking. Bin size 10 ms. **(D)** Schematic of the first photostimulation frame of the bar-like photoactivation on the map of S1 barrels. P: posterior, M: medial. **(E)** Snapshots of the cortical surface illustrating bar-like photostimulation frames for each virtual cursor position. Only when the virtual cursor was in position 5, licks were rewarded.

After initial sessions where we habituated the mice to remain head-fixed and lick for water, we trained the mice to solve a 1D cursor control task via the brain-machine interface. To this aim, we sorted 3 “Master” neurons from the raw M1 neuronal activity (7 Master neurons in two mice, see Methods). The activity of these neurons controlled the movements of a virtual cursor during the sessions. Their summed firing rate was measured every 10 ms and was smoothed with a 100 ms kernel. It was then normalized by the firing rate distribution measured during a 3-minute baseline at the start of each session. Finally, we discretized the normalized values into 8 positions of a virtual cursor (see Methods, Fig. 1C and Fig. S3).

Whenever the virtual cursor was in the rewardable position (Fig. 1C-E, only in position 5, except in our first experiments — see Methods), the mice could obtain a water droplet by licking a port located next to their tongue. Water rewards could be triggered only by licking on the spout. Therefore, in the absence of licking while the virtual cursor was in the rewardable position, no water was made available to the mouse. During the task, the current position of the virtual cursor was provided online to the mice through patterned optogenetic stimulation of S1 that triggered local, low-latency spiking activity (see Methods). The mice expressed constitutively channelrhodopsin in pyramidal neurons (Emx-Cre;Ai27 strain, (*31*). The photostimulations were dynamically updated, with an intrinsic hardware latency of 12±5 ms from the firing of the Master neurons to the corresponding photostimulation update (*15*).

At each time point, the pattern of cortical illumination consisted of focused spots that targeted five of the S1 barrels. We arranged these spots to form a bar-like arrangement of barrel activations, sweeping on barrels corresponding to caudal whiskers for position 1 of the virtual cursor, up to rostral whiskers for position 8 (Bar feedback, Fig. 1D,E).

A 30 min training session per day was delivered during five days. To obtain more rewards, mice had to increase the amount of time during which the virtual cursor was in the rewardable position, and/or improve their ability to lick in those time windows.

When Bar feedback was provided (Fig. 2A, top), the mice were able to increase significantly their performance within the 5 consecutive training sessions (example in Fig. 2B, top). On the first session the mice licked occasionally but the virtual cursor was almost never in the rewardable position at the same time, and the mice obtained almost no water. On the fifth day training session, the same mice performed licking bouts at times when the virtual cursor entered the rewardable position, and thus obtained rewards more frequently. Overall, over the course of five training sessions, the performance measured as the frequency of rewards (licks per seconds) significantly increased more than ten times (from 0.014 to 0.19 rewards/s; orange curve of Fig. 2C, Mann-Whitney U = 5, p < 0.001, n = 10 mice). In contrast, in the absence of optogenetic feedback (Fig. 2A,B, bottom), the mice failed to reliably increase the frequency of rewards despite the same amount of training (0.025 vs 0.022 rewards/s; gray curve in Fig. 2C, Mann-Whitney U = 31, p = 0.48, n = 8 mice among the 10 tested in the Bar feedback condition).

**Fig. 2.**
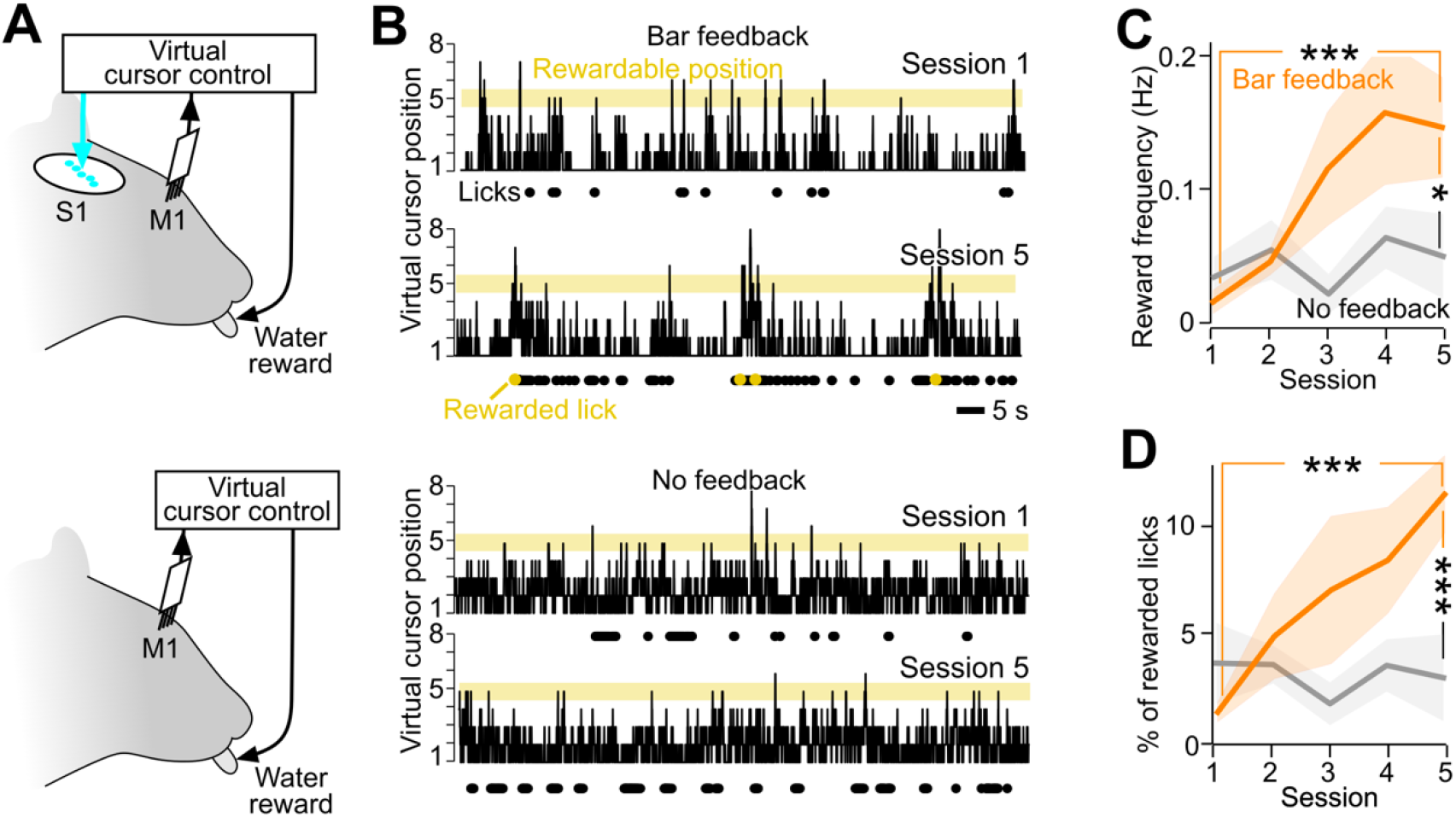
Sensory feedback to the whisker part of S1 enhances task performance. **(A)** Schematic of the Bar feedback and No feedback conditions. **(B)** Position of the virtual cursor computed from the merged activity of the Master neurons, in the first versus the fifth training session of one mouse, in the Bar feedback condition (top) and in the No feedback condition (bottom) (100 s displayed). Yellow background: rewardable position. Black dots: lick times. Yellow dots: rewarded lick times. **(C)** Performance quantified by the average frequency of rewards per session across training, comparing the Bar feedback condition (orange, 10 mice) and the No feedback condition (grey, 8 mice). Shaded backgrounds: ± standard error of the mean (SEM). *: p < 0.05. **: p < 0.01. ***: p < 0.001, non-parametric Mann-Whitney tests. **(D)** Same as C for the specificity of licking, quantified as the proportion of rewarded licks among all licks, across behavioral sessions.

Importantly, the increased reward frequency in the Bar feedback condition was accompanied by an increase in the specificity of licking, measured by the percentage of licks that were rewarded among all licks (Fig. 2D, Mann-Whitney U = 1, p < 0.001). This indicated that the mice did not simply increase their licking frequency irrespective of the virtual cursor position in order to solve the task. We conclude from these data that the optogenetic feedback to the barrel cortex was required for learning to control this brain-machine interface within five training sessions.

### Shuffling the spatiotemporal structure of the feedback disrupts learning

We hypothesized that in these initial experiments, the specific spatio-temporal structure of the Bar feedback helped the mice to control the virtual cursor, whereas other types of feedback might not result in similar fast task learning. To explore this question, we selected a subset of 3 additional feedback conditions that degraded the spatio-temporal structure of the original Bar feedback in controlled ways (Fig. 3A).

**Fig. 3.**
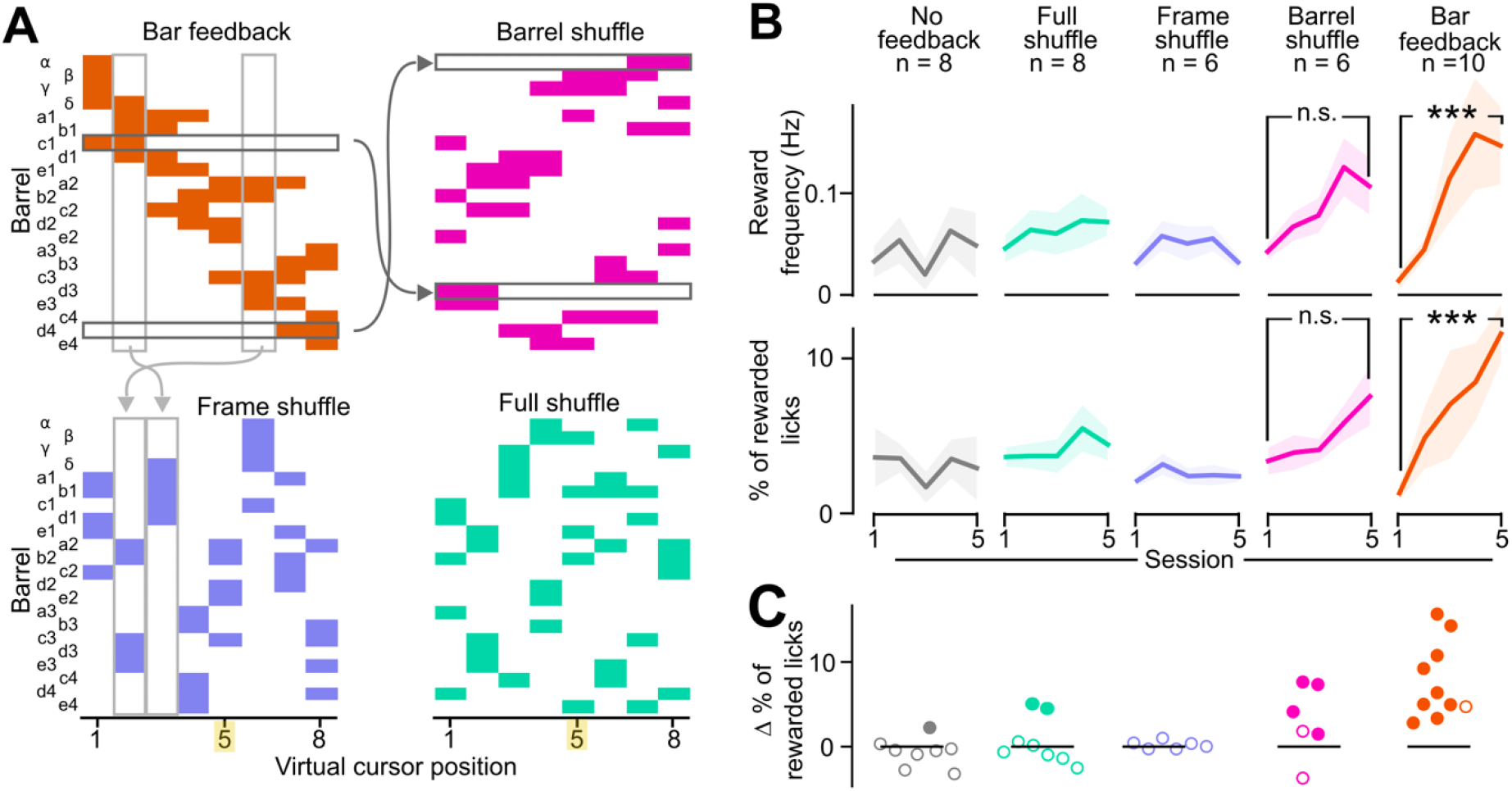
Disrupting the spatiotemporal structure of the bar feedback impairs learning. **(A)** Spatial and temporal structure of the feedback across frames in the four tested conditions. Horizontal arrows: barrel identity permutation to generate the Barrel shuffle from the Bar feedback. Vertical arrows: frame identity permutation to generate the Frame shuffle from the Bar feedback. Yellow highlight: rewardable virtual cursor position. **(B)** Reward frequency (top) and percentage of rewarded licks (bottom) of the mice over 5 training sessions. ***: p < 0.001, non-parametric Mann-Whitney tests. Shaded backgrounds: ± standard error of the mean (SEM). Bar feedback and No feedback data is the same as in Fig. 2. **(C)** Difference between the proportion of rewarded licks of the mice between the first versus fifth training session. Each point represents a mouse. Filled point: bootstrap significance test, p<0.05.

In the *Barrel shuffle* condition, we degraded the spatial arrangement of the Bar feedback by randomly shuffling the identity of the photostimulated barrels, therefore removing the contiguity and spatial alignment between simultaneously activated barrels, but preserving temporal overlap of 2-3 barrels from one frame to the next (6 mice; see all photostimulation frames in Fig. S4). In the *Frame shuffle* condition, we preserved the spatial organization of the photoactivated barrels within one frame, while in contrast, the correspondence of the frames with the virtual cursor position was shuffled. This rearrangement disrupted the overlap and contiguity of the displayed frames during evolutions of the virtual cursor (6 mice). Finally, in the *Full shuffle*, both the spatial position of the barrels and the frame-to-cursor correspondence were randomized (8 mice).

We trained mice to control the virtual cursor by M1 neuronal activity while receiving these different feedback patterns. Apart from the spatial content of the optogenetic frames themselves, training was identical to that implemented for the Bar feedback and No feedback conditions. In these experiments, the mice remained actively engaged. They licked and obtained rewards throughout all training sessions (Fig. S5A). However, in contrast to the Bar feedback condition, we found no significant increase in mice performance across sessions (Fig. 3B). In the Barrel shuffle condition, we noticed a trend towards an increase in the reward frequency and in the percentage of rewarded licks, but it did not reach significance (Fig. 3B, reward frequency: Mann-Whitney p = 0.064; % rewarded licks p = 0.132) although 4 mice out of 6 did show a significant increase in the percentage of rewarded licks (Fig. 3C).

Through these experiments, we consecutively trained mice to learn the task with multiple different feedback structures. Therefore, the order of the training sequence might have had an impact on the learning performance. We explored this potential effect with two groups of three mice which followed consecutively training in the No feedback, Full shuffle and Bar feedback conditions, in two different orders (Fig. S5B). Irrespective of the protocol training order, significant learning was observed only in the Bar feedback condition (Mann-Whitney U = 0, p = 0.04). We conclude that the order of the different training protocols did not impact the learning process. Overall, we found that the spatiotemporal structure of the feedback impacted heavily the behavioral performance of the mice, and that the Bar feedback enabled fastest learning.

### Mice learn to lick during time windows of reward availability

Next, we asked which mechanisms could underlie the ability of mice to improve their performance over 5 sessions. One hypothesis is that the mice improved their licking behavior by timing their licks more accurately relative to entries of the virtual cursor in the rewardable position. Given the tendency of mice to lick in long rhythmic bursts, we focused on the onsets of licking bouts, with the assumption that they indicate the attempts of the mice to obtain reward (Fig. 4A). We computed the proportion of lick burst onsets that fell within ± 100 ms of the virtual cursor entry in the rewardable position, which is approximately the duration of a tongue licking cycle. We found that this proportion increased significantly only in the Bar feedback condition (Fig. 4B,C; Mann-Whitney p < 0.01), and that again, a similar, non-significant trend was visible in the Barrel shuffle condition. We conclude from this data that the mice did learn to adjust their licking patterns to the virtual cursor dynamics.

**Fig. 4.**
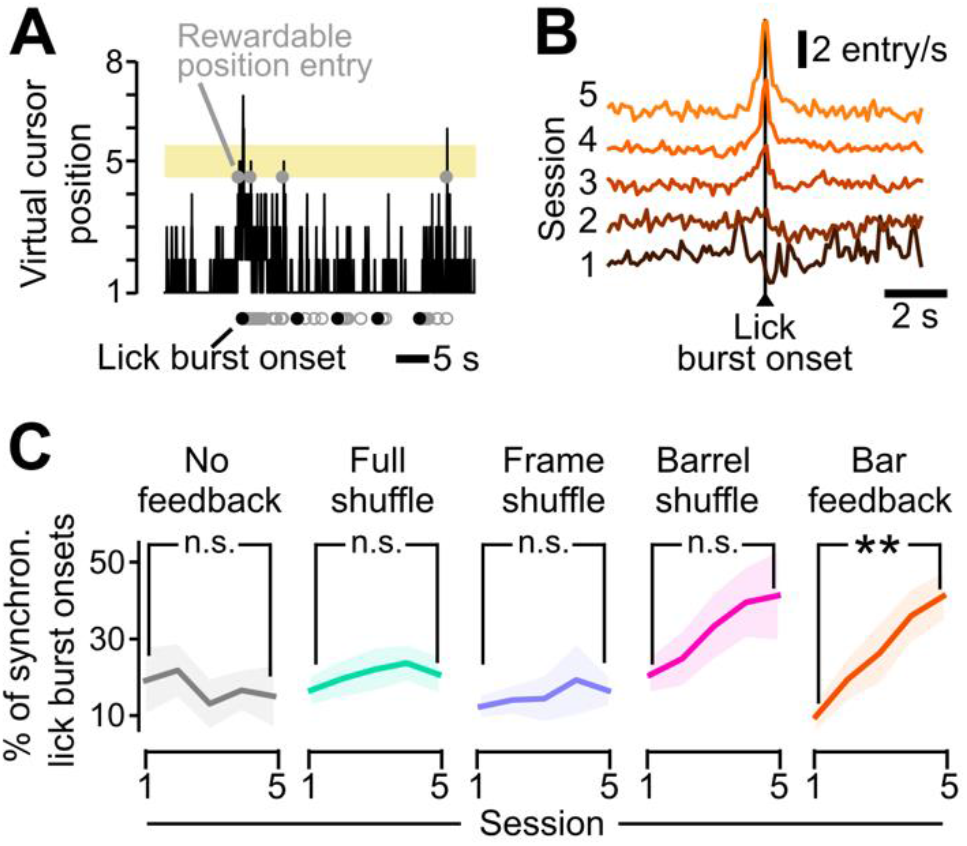
Mice learned to synchronize licking with the entries of the virtual cursor in the rewardable position. **(A)** Example of the virtual cursor position as a function of time. Gray open circles: licks. Black dots: onsets of lick bursts. Gray dots on virtual cursor position 5 indicate entries of the virtual cursor in the rewardable position. To avoid confusion, rewarded licks are not highlighted. **(B)** Population average time histograms of the number of entries of the virtual cursor in the rewardable position around all lick burst onsets, for the Bar feedback condition, across the five training sessions. Baseline levels were shifted upward for clarity. **(C)** Percentage of lick bursts that are synchronous (within ± 100 ms) with entry of the virtual cursor in the rewardable position. **: Mann-Whitney, p < 0.01. Shaded backgrounds: ± SEM.

### Mice learn to bring the virtual cursor in the rewardable position dynamically

In parallel to this adaptation of the licking behavior, there could also be a change in the dynamics of the virtual cursor near and in the rewardable position, providing more opportunities for rewarded licks. To explore this hypothesis, we analyzed the virtual cursor dynamics at different time scales, focusing on how it changes from the first to the fifth training session. First, we measured the average time spent in the rewardable position across the whole duration of a session (Fig. 5A). We found that it increased significantly in the Bar feedback condition (Mann-Whitney p < 0.05), in contrast to all other tested feedback conditions. This confirms that mice learned to bring the cursor in the rewardable position more often. When we plotted the average virtual cursor position in time, first on a long timescale, we noticed that the curves for Sessions 1 and 5 started at the same level, followed by an upward shift 10 to 15 s after the start of the photostimulation in Session 5 (Fig. 5B; cursor position significantly higher in 15-100 s vs. 0-10 s, Wilcoxon test p = 0.014, only for the Bar feedback condition).

**Fig. 5.**
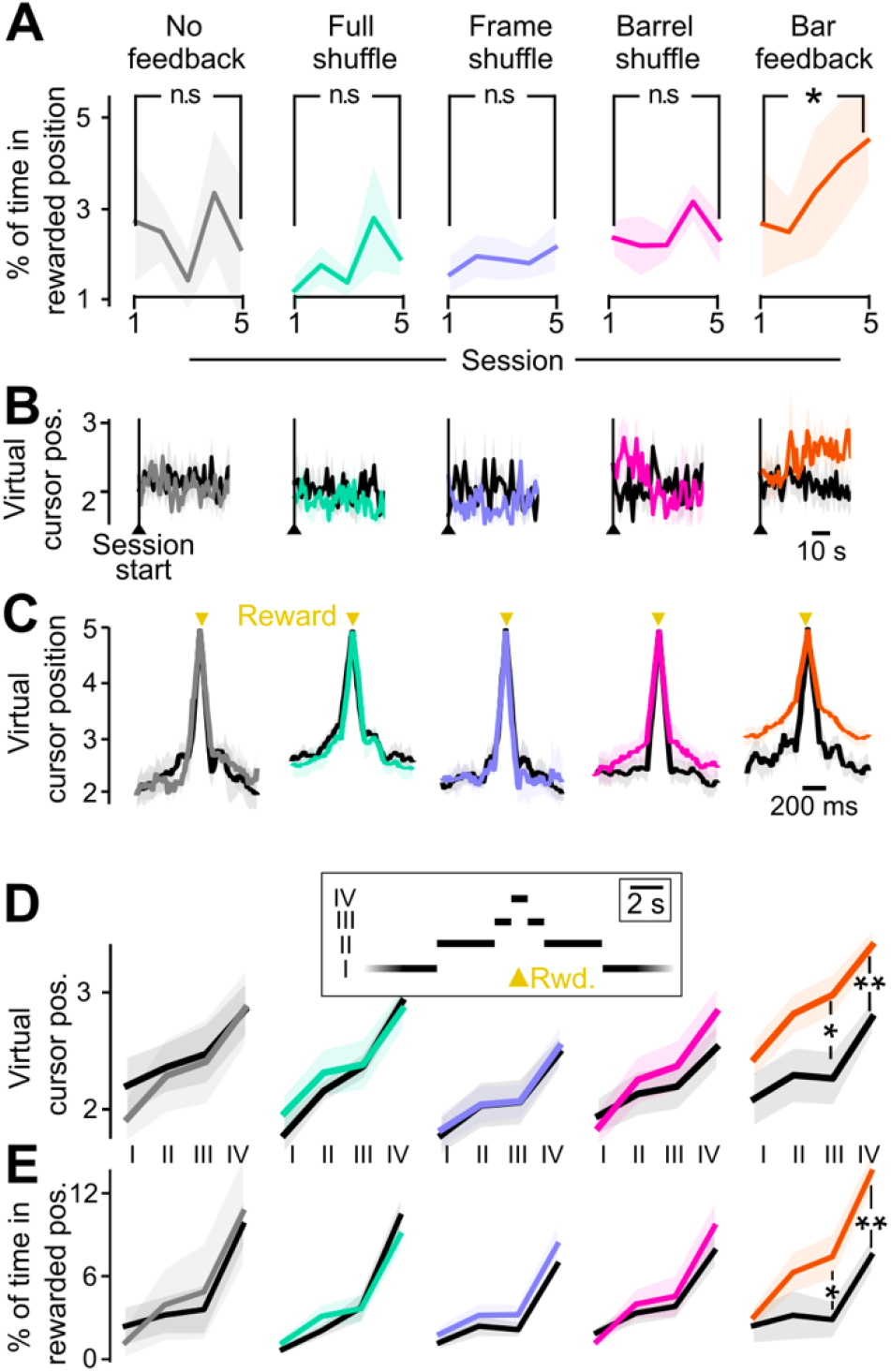
Bar feedback enables the mice to actively control the virtual cursor position so that they spend more time in the rewardable position. **(A)** Proportion of time spent in the rewardable virtual cursor position (position 5) over the whole session duration. *: Mann-Whitney, p < 0.05. Shaded backgrounds: ± SEM across mice. **(B)** Average virtual cursor position ± SEM at the onset of the session, in the first versus last training session. Vertical line: start of the session, which is also the start of the photostimulation. **(C)** Average virtual cursor trajectory, aligned to the reward times, in the 5 feedback conditions. Black: first session. Colors: session 5. Shaded backgrounds: ± SEM across mice. **(D)** Average virtual cursor position, in four time windows around reward: (I) More than 5 s away from any reward. (II) 1.5 to 5 s away. (III) 0.5 to 1.5 s away (IV) Within 0.5 s of a reward. *: Mann-Whitney, p < 0.05. **: p < 0.01. **(E)** Percentage of time spent in the rewardable position, in the four time windows around reward defined in d. Mean ± SEM across mice.

These delayed dynamics rule out the hypothesis that photostimulation could have non-specifically increased the overall activity, and consequently the virtual cursor position. We then investigated whether there was a dynamical control of the virtual cursor on a faster time scale leading to rewards. We observed that in the Bar feedback condition, and only in this condition, the mean cursor position was significantly larger after training than on session 1, up to 1.5 s around reward occurrence (Fig. 5C,D, Mann-Whitney, p < 0.01). In the same time windows, the virtual cursor spent a proportion of time in the rewardable position that was significantly larger after training compared to before (Fig. 5E, Mann-Whitney, p < 0.01). There were no changes in these measures in epochs further away from rewards (1.5-5 s and > 5 s from any reward, Mann-Whitney, p > 0.05), indicating that there was not a systematic additive shift in the virtual cursor position throughout the session, but rather numerous short explorations of higher cursor positions around rewards.

Overall, these results suggest that during training, the mice learned to manipulate the virtual cursor and bring it in the rewardable position more often, thus creating more opportunities for enhancing their performance by well-timed licks.

### One Master neuron dominates control of the virtual cursor

By design, changes in the dynamics of the virtual cursor are a direct consequence of changes in the underlying Master neurons activity, albeit in a non-linear way tailored to each mouse extracellular recording (see Methods and Fig. S3). We verified the changes in firing rate underlying the observed changes in virtual cursor trajectory. In particular, since the activity of several Master neurons was summed to drive the cursor, we wondered whether all Master neurons contributed equally, or if instead motor control of the virtual cursor was dominated by a subset of the Master neurons. To investigate this question, we sorted the Master neurons as a function of their contribution to the virtual cursor position at reward time, and we looked at the evolution of their spiking activity over training. We termed “dominant” the Master neuron that on average fired the most at reward time, in a +/- 100 ms window.

First, we checked the firing rate of Master neurons at the time scale of a whole session. Importantly, right at photostimulation onset, there was no detectable change of activity of Master neurons (Fig. 6A). This indicates the absence of an immediate photostimulation effect, in agreement with what we had observed on the virtual cursor position (Fig. 5B). Interestingly, the dominant Master neuron had a markedly increased firing rate in Session 5 compared to Session 1, including in the baseline period before photostimulation start (Fig. 6A). When averaged across the whole duration of each session, the firing rate of the dominant Master neuron was indeed significantly larger after training compared to before, and larger than any non-dominant Master neurons (Fig. 6B, Mann-Whitney, p < 0.01, see also Fig. S6A). This increase was specific to the Bar feedback condition. Non-dominant neurons, on the contrary, showed little change in activity after training, in all feedback conditions (Fig. 6A,B).

**Fig. 6.**
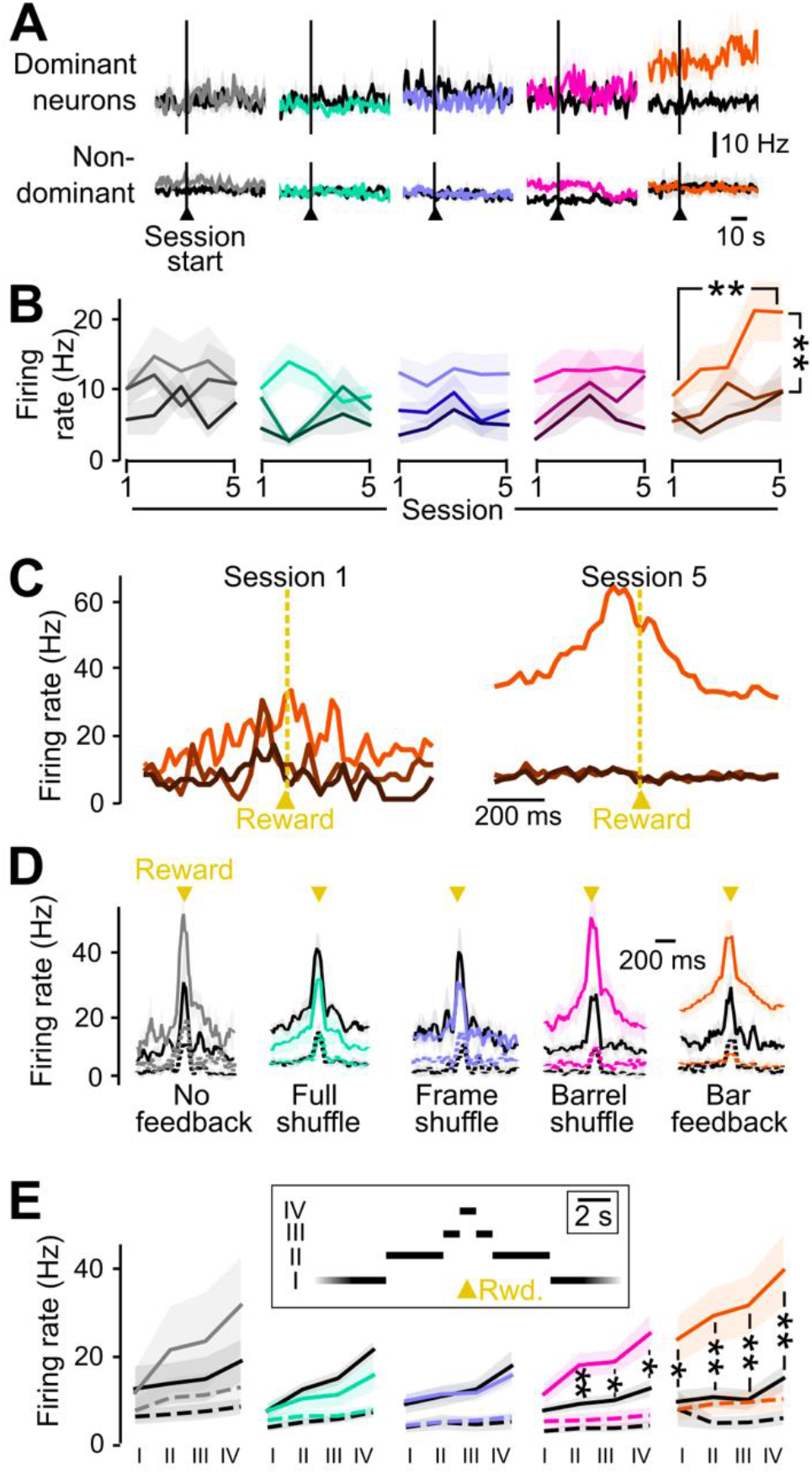
Emergence of a dominant Master neuron in the Bar feedback condition. **(A)** Firing rate at the onset of the session, for dominant (top) and non-dominant (bottom) Master neurons in the first (black) versus last training session (colors). Shaded backgrounds: ± SEM across mice. **(B)** Mean firing rate of the dominant (saturated colors) and non-dominant Master neurons, over training sessions. Shaded backgrounds: ± SEM across mice. **: Mann-Whitney, p< 0.01. **(C)** Mouse case study of the time histogram of the activity of Master neurons around rewards, in the Bar feedback condition, sorted from the weakest (dark brown) to the dominant neuron (saturated orange) at the time of the reward, in the first (left) versus the fifth (right) training sessions. **(D)** Time histogram of the activity of Master neurons around rewards, in the 5 tested feedback conditions. Session 1 is shown in black, and session 5 in saturated colors. Continuous line: dominant Master neuron. Dashed line: average of non-dominant neurons. **(E)** Average firing rate of dominant (continuous line) and non-dominant (dashed line) Master neurons in the first (black) versus the last training session (colors), measured in the same time windows as in Fig. 5: (I) More than 5 s away from any reward. (II) 1.5 to 5 s away. (III) 0.5 to 1.5 s away (IV) Within 0.5 s of a reward. Shaded backgrounds: ± SEM across mice. *: Comparison between the first and fifth training session. Mann-Whitney p value < 0.05; **: p < 0.01.

These observations on a long timescale could be due to persistent elevated firing after training, or to an accumulation of short bursts of high firing of the dominant Master neuron. We thus analyzed the fast dynamics around rewards, as done previously for the virtual cursor position (Fig. 5C). Figure 6C shows the firing rate of individual Master neurons around reward times, for Sessions 1 and 5 of one mouse trained in the Bar feedback condition. After training, one of the Master neurons showed a much higher firing rate with a prominent peak around the reward time. Population averages across mice confirms this tendency for the dominant neuron in the pool of Master neurons, whereas little changes were observed on non-dominant neurons (Fig. 6D). Again, this was specific to the Bar feedback condition, although a more moderate trend was noted for the Barrel shuffle condition. We quantified the firing rate in several time windows around rewards (similar to Fig. 5D for the virtual cursor position). The firing rate of dominant Master neurons around reward times in the Bar feedback condition showed a strong and significant increase after training compared to before (Fig. 6E). This increase was specific to the dominant Master neurons, and was highest around reward times (Mann-Whitney p < 0.01). It was less pronounced but still significant more than 5 s away from any reward (Mann-Whitney p = 0.04), an observation that we relate to the elevated firing rate in the baseline period already before the task started (Fig. 6A). We observed a similar, but more limited phenomenon in the Barrel shuffle condition (Fig. 6E).

Note that the tonic component of the increased firing rate of the dominant Master neuron was normalized by the motor control algorithm, and it could therefore not lead to a shift in virtual cursor position (see Methods). Thus, we conclude that in the Bar feedback, and to a lesser extent in the Barrel shuffle, the mice learned to control the virtual cursor position mostly by increasing the activity of one Master neuron in short bursts of elevated firing around lick times, enabling them to obtain rewards.

### Playback experiments confirm the role of active motor control for task performance

Finally, to further explore the role of motor control on task performance, we performed playback experiments on three mice that had already learnt the full closed-loop task with the Bar feedback protocol. The mice received the same optogenetic stimulation sequence as in their last closed-loop session with Bar feedback, and they could still receive reward by licking when the virtual cursor was in the rewardable position. However, the virtual cursor dynamics was now independent from the ongoing activity of motor cortex neurons. In other words, the animals were relieved of the motor control aspect of the full task (Fig. 7A,B). Interestingly, the frequency of rewards dropped significantly for each mouse in the playback condition (Fig. 7C) even though by design, the virtual cursor spent as much time in the rewardable position as during the Bar feedback last session. Analysis of the synchrony between licking onsets and the entries of the virtual cursor in the rewardable position revealed that these events were not coordinated anymore (Fig. 7D).

**Fig. 7.**
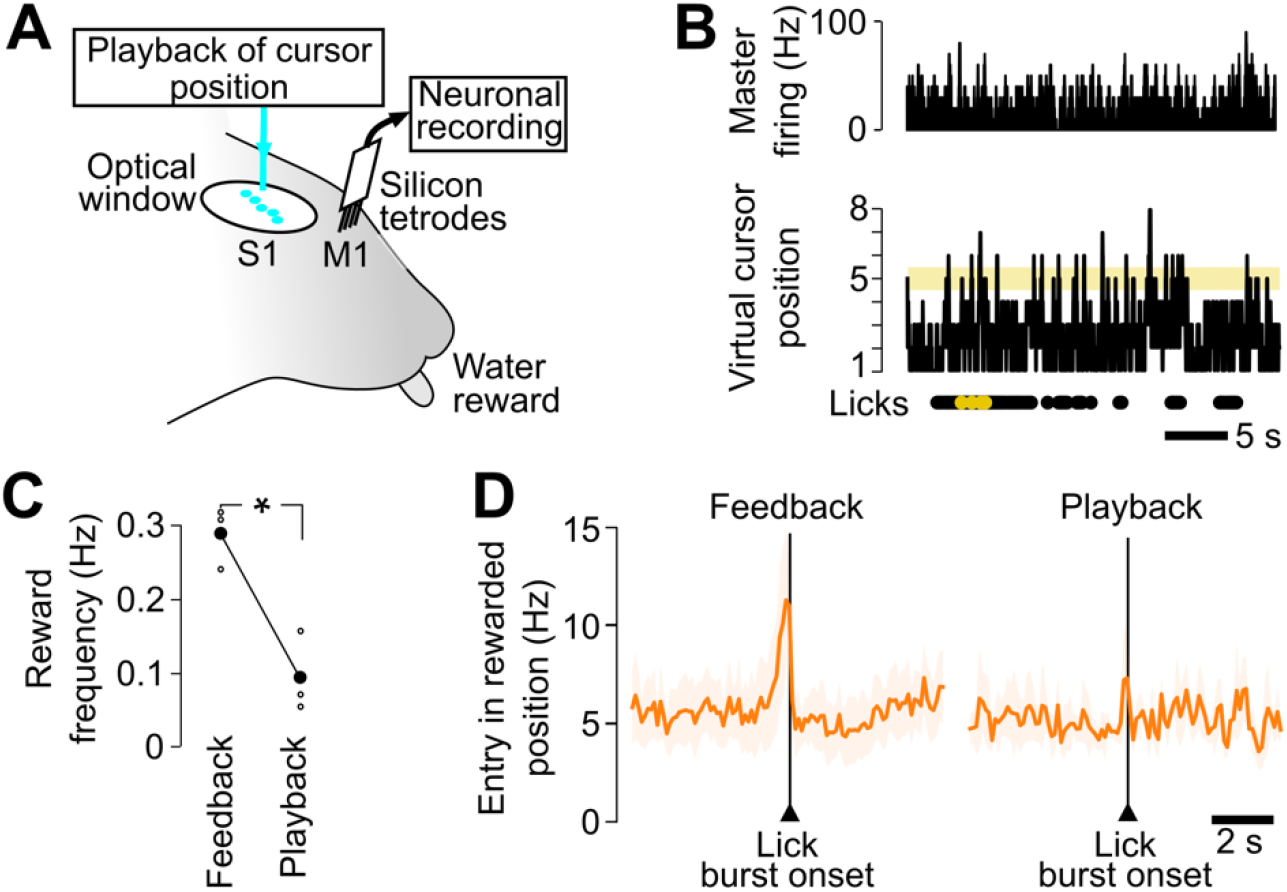
Lick timing is not accurate in a playback condition. **(A)** Playback configuration with chronic extracellular recording in M1 and Bar feedback optogenetic stimulation on barrels in S1. Previously acquired sequences of cursor positions are played back, independent from M1 firing rates. As in closed-loop sessions, reward delivery is contingent on synchronous 1) licking and 2) presence of the virtual cursor in the rewardable position. **(B)** Top, histogram of Master neurons activity during a playback session (30 seconds shown). Bottom, time course of the virtual cursor position, disconnected from the Master firing. Below: licks, and rewarded licks during the same interval. Bin size 10 ms. **(C)** Frequency of rewards during the last session with closed-loop Bar feedback and the session with open-loop playback. *: Mann-Whitney, p < 0.05. Gray background: SEM. n = 3 mice. **(D)** Histogram of lick bursts onsets, with respect to the times of entry of the virtual cursor in the rewardable position around the onset of lick bursts, for the last session with closed-loop Bar feedback (left) versus the session with open-loop Bar playback (right), averaged for the three tested mice.

To confirm that active motor control is necessary not only for task execution but also for learning, we trained three naive mice to perform the playback task during 5 sessions. Consistent with the previous playback result, we found that the mice failed to increase significantly their performance during this playback training (Fig. S7).

Overall, these playback experiments demonstrate that in the Bar feedback condition, the mice did not only respond to sensory cortex stimuli by licking, but instead actively coordinated their licking with timely modulations of the cursor position.

## Discussion

In this study, we demonstrate that in the context of improving motor control of a brain-machine interface, the integration of direct cortical feedback can be heavily impacted by its spatiotemporal organization. Specifically, we trained mice in a task for which the brain-machine interface could be used to move a virtual cursor into a rewardable zone. We found that the performance after training was highest when feedback provided the position of the cursor in the form of a bar-like photostimulation across the cortical surface (Bar feedback condition). In contrast, we found that when we disrupted the spatial contiguity of simultaneously stimulated barrels (Barrel shuffle), learning was clearly reduced, and when we disrupted the continuity of the bar in time (Frame shuffle), it went down to levels observed without feedback.

This difference in performance was associated with a reorganization of the ongoing neuronal activity that was specific to the Bar feedback condition. More precisely, one of the M1 Master neurons driving the cursor became dominant in terms of activity levels and led the virtual cursor to spend more time in the rewardable position, thereby increasing the opportunity for rewards. In parallel, licks were more synchronized with entries in the rewarded position.

### A fast bidirectional BMI setup for the mouse

Current research aimed at integrating somatosensory feedback in a cortical brain-machine interface relies on invasive techniques of recording and stimulation in awake behaving animals. Pioneering teams are developing prototypes in non-human primates as well as human participants (*9*, *32*). Here, we have developed a novel brain-machine interface tailored to the mouse whisker system, a sensorimotor loop that has been described in a comprehensive way, from the cellular to the network level (*21*, *22*). This approach has allowed us to take advantage of recent optogenetic tools available for these animals. We could activate excitatory neurons in the cortex according to spatial light patterns that were adapted, in each individual mouse, to the topographic map of the whiskers present in S1. Furthermore, we benefited from our low-latency (12 ± 5 ms) closed-loop design which enables the delivery of feedback in a dynamic way, so that the ongoing activity of the Master neurons controlled online the stimulation frames. Indeed, a low-latency somatosensory feedback could be an important parameter in the context of sensorimotor learning (*33*).

To provide distributed feedback to the mice, we chose to generate illumination patches above individual S1 barrels, rather than try to mimic the broad spread of activity that is generated by multiple whisker stimulation sequences (*27*). The rationale has been to mimic activation patterns of multiple lemniscal thalamic inputs, which are known to project into barrel columns of corresponding whiskers, and which should then trigger broader activation of the cortex through intracortical connectivity, both within and across layers (*22*). We hypothesize that this recruitment of intracortical mechanisms is key to the similarity between artificial and physiological stimulation. We certainly acknowledge that significant differences remain between the optogenetic activation and physiological activation of the barrel cortex. In particular, we did not attempt to reproduce non-lemniscal thalamic input patterns which don’t follow a clear topographical mapping at the surface of the cortex, and which are thus difficult to activate specifically.

### Impact of somatosensory feedback on neuroprosthetic learning

In our experiments, in the absence of optogenetic feedback, the mice failed to learn the task. In contrast, a few previous studies have suggested that BMI learning could take place without any feedback of the conditioned neuronal activity to the animal (*34*, *35*) Several differences could explain these seemingly opposite results. First, in those studies, the animals received the reward automatically once the neuronal activity reached the predefined threshold. In contrast, in our task, the mice have to learn also to lick in order to obtain the reward. This combination of firing rate modulation and required licking probably makes the task much more challenging. Second, in our study, movements of the virtual cursor occurred on average every 50 ms, so that temporal precision of licking was important. This must also have been challenging, particularly in the absence of any feedback. These reasons could explain the lack of learning that we report in the No feedback condition.

Our study shows that direct cortical feedback can enable the learning of a sensorimotor task in these conditions, pending that feedback with an adequate spatio-temporal structure is provided. This is consistent with recent work exploring cortical somatosensory closed-loop BMIs in humans with intracortical electrical stimulations (*8*), as well as with previous work emphasizing the prevalent role of ongoing sensory feedback in motor learning (*33*, *36*).

Importantly, our experimental design did not incorporate a physical implementation of a device to be moved by the animal towards a target. Instead, we computed the position of a virtual cursor and used it to select the next frame of the ongoing feedback. This choice ensured that the optogenetic feedback delivered to S1 was the sole source of sensory information about the virtual cursor position available to the animal during the task. This is in contrast to most previous closed-loop BMI studies, in which ongoing visual feedback of the neuroprosthesis was always available for adjusting motor control in addition to cortical stimulation (*9*, *32*).

### Motor control of the virtual cursor

In this study, direct demonstration of voluntary motor control was challenging because virtual cursor movements were generated continuously rather than triggered. Still, we found several indications of active motor control of the virtual cursor, that were specific to the Bar feedback condition, and to a lesser extend to the Barrel shuffle condition. In particular, only in this feedback condition did the virtual cursor position shift towards the position of the rewarded frame, as the mouse prepared to collect rewards in the next seconds (Fig.5D,E). In addition, analysis of the neuronal activity of the Master neurons underlying the virtual cursor position revealed that throughout learning sessions, neuronal activity evolved towards the dominance of a single one of their Master neurons, in particular during the modulations of activity towards the rewarded position. This rearrangement took place only in the Bar feedback condition (Fig. 6). Finally, during additional playback sessions at the end of a sequence of training in the Bar feedback condition, the mice appeared unable to maintain the performance level they attained during previous closed-loop training sessions, indicating that active motor control was required for performance (Fig. 7).

Overall, we conclude from these analyses that in the Bar feedback condition, the mice did rely on the active modulation of their Master neurons to collect rewards. The lack of such motor control in other feedback conditions illustrates the impact of the spatiotemporal structure of our distributed feedback, not only for sensory information processing, but more generally for sensori-motor integration of the feedback.

Regarding the playback experiments, we should point out that in one study, after operant conditioning of motor cortex neurons based on a single barrel S1 optogenetic feedback, mice were able to efficiently gather rewards during playback training (*16*). Similarly, we have previously shown that in our experimental setting, mice were able to detect a static, single frame of the Bar feedback to obtain rewards (*15*), Fig. 5) or to track a continuous, slowly rotating bar (*37*). We hypothesize that what makes the playback condition here (Fig. 7) comparatively more challenging than these previous experiments is that it combined rapidly changing feedback with a distributed, more complex spatial pattern. In addition, a low-latency licking was necessary when the cursor entered the rewardable frame. All these challenges meant that to be successful, the mice had to anticipate the entrance in the rewardable frame, as the cursor could escape the rewarded position within milliseconds. In contrast to the playback condition, we hypothesize that in the closed-loop Bar feedback condition, the motor control of the virtual cursor provided the degree of rewarded frame anticipation that allowed timely licks and an increase in the proportion of rewarded licks.

### Pattern contiguity impacts learning and performance

We showed that direct cortical feedback should obey spatiotemporal rules of organization in order to be efficiently integrated into motor control. Specifically, we found that the mice were able to learn to control a virtual cursor using an S1 bar-like feedback that featured contiguity of the activated barrels both within a given frame, and across consecutive frames.

When across-frames contiguity was removed, in the Frame shuffle feedback condition and also the Full shuffle condition, we found no sign of learning, as in the absence of feedback altogether. We hypothesize that the lack of temporal continuity across consecutive feedback frames may have prevented the anticipation of upcoming cursor movements. Given our fast-paced cursor positioning task, this translated in an inability to learn the task. This hypothesis is consistent with the findings in our previous, open-loop work (*37*).

However, when only within-frame contiguity was removed (Barrel shuffle condition) learning was at intermediate levels. The mice were able to exploit the feedback to some degree, but lacked the accuracy that is required to really synchronize virtual cursor and licking efficiently. These results on the relevance of both the spatial and the temporal structure of intracortical feedback suggest that the sensorimotor task of driving the virtual cursor to the target draws upon pre-existing features of S1-M1 microcircuits, linked to their topographic organization (*38*). When the contiguity of the feedback was disrupted, the functional architecture of the cortex may not have been adapted anymore to the novel sensorimotor computations that were required to solve the task. Thus, learning to extract the relevant virtual cursor information from the different shuffled conditions may require multiple additional training sessions, if indeed the required functional connections can be recruited from the existing anatomical scaffold (*39*). In fact, previous work does suggest that learning a spatially shuffled cortical stimulation is possible if training spans multiple training sessions, with the assistance of visual feedback (*19*, *40*). This seems consistent with the signs of learning that we did observe in the Barrel shuffle condition (Fig. 3B,C).

Interestingly, in contrast to the limited capacity for processing sensory feedback with different feedback structures, we observed that the animals were able to readily adapt to the constraints that were set on the motor side. Indeed, consistent with previous studies (*16*, *30*, *41*, *42*), we found that an arbitrary set of a few M1 neurons could be conditioned in an operant way to learn to control a virtual cursor along one dimension. The sharp contrast between the necessity of spatially structured patterns on the sensory side, and the adaptability of the neuronal networks on the motor side, could have several explanations. One is that we used mesoscale patterns to encode sensory feedback, encompassing large numbers of neurons and connections. Plastic reorganization at this scale may be more demanding than when targeting only a few neurons. Indeed, there is evidence that as the number of neurons controlling motor brain-machine interfaces increases, it becomes necessary to take into account their initial functional connections in order to learn to control the prosthesis rapidly (*43*-*45*). Alternatively, another possibility is that primary sensory cortical circuits may be intrinsically less plastic than motor ones during motor skill learning (*46*). Future experiments will need to address this question.

### Functional role of the S1 somatotopic map

So far, the contribution of cortical maps to sensory information processing in general has remained unclear (*47*) despite the thorough descriptions of the maps in primary sensory cortices. In the case of the barrel cortex, several of the functional properties encoded by its neurons are spatially organized inside the map beyond spatial topography (*48*-*51*), including some multi-whisker features (*27*, *49*). The large-scale organization of feature encoding would be favored because of the dense lateral connectivity inside S1, enabling distributed cortical computations (*24*). Through this rich anatomical substrate, non-linear spatiotemporal integration in S1 results in enhanced responses to some input patterns, and suppression of responses to other patterns (*52*). However, so far, these feature extraction properties have not been causally linked to behavior, except the somatotopy itself recently (*37*).

Our results shed light on the functional role of topography of the somatosensory cortical map in the behaving animal, by testing causally the impact of different patterns of sensory input. In particular, our work reveals that feedback patterns that are contiguous within the frame of the barrel cortex topographical organization are best suited to sensorimotor integration. Such optimal patterning of dynamical distributed feedback could be combined with other means of transmitting feedback information to the brain, such as temporal and amplitude modulation of stimulation pulses (*14*, *16*, *53*).

Finally, current BMI prototypes require long training and lack precision and flexibility, probably because they lack the appropriate somatosensory feedback (*54*). From our results, we propose that feedback strategies based on intracortical stimulation should favor spatial and temporal continuity within the known topography of the target areas. We hope that unveiling such fundamental constraints of neuronal circuits will enable the development of a new generation of BMIs, incorporating rich proprioceptive and tactile feedback essential to achieve dexterity and embodiment.

## Materials and Methods

### Mouse preparation

All animal experiments were performed according to European and French law as well as CNRS guidelines and were approved by the French ministry for research (ethical committee 59, authorization 858-2015060516116339v5 and 25932-2020060813556163v7). The data were obtained from 16 adult (6-10 weeks old) Emx1-Cre;Ai27 mice (*31*). The brain-machine interface methodology has been published previously (*15*). All surgeries were performed under isoflurane anesthesia in 100% air. Isoflurane concentration was adjusted in the range 1–4% depending on mouse state, assessed by breathing rate and response to tail pinch. Each mouse underwent two surgeries. During a first surgery, a 5 mm diameter glass optical window was implanted over the left primary somatosensory cortex (S1, P-1.5 mm and L3.3 mm from bregma, (*55*) and a head-fixation bar was implanted on the contralateral side of the skull (*56*). Eight days later, the clarity of the optical window was assessed, and if adequate, intrinsic imaging was performed to locate the S1 barrels (see below). If this first step was successful, a second surgery was performed to chronically implant (*57*) a 32 channel silicon probe in the shape of 8 tetrodes (A1×32Poly35mm25s 177A32, NeuroNexus, USA, Fig. 1a-b, Fig. S1,S2). The electrode was implanted in the whisker zone of the motor cortex (M1, A1.5 mm L0.6 mm from bregma, electrode recording sites 650 – 800 μm deep in cortex).

### Chronic neuronal recordings

Following the second surgery, mice were monitored for 5 days to allow the extracellular recordings to stabilize (bandpass 1Hz - 6000Hz). We then characterized the shape and amplitude of the units isolated by the online spike sorting (Blackrock microsystems, USA). Clusters corresponding to well-defined single units (consistent spike shape and an adequate autocorrelogram, with a clear refractory period, see Fig. 1b) were manually selected within the tetrode spike amplitude space. This manual selection was controlled before each session to ensure that we maintained unit separation while keeping track of the same units across sessions (Fig. S1). Once the online spike sorting was ready, the training session begun. At the start of the training sessions, we recorded a median of 25.5 neurons simultaneously, with an interquartile range (IQR) of 5.25 neurons (n = 10 mice). After 17 days (average last training session) we recorded a median of 25 neurons (IQR = 16 neurons, n = 10 mice).

### Brain-machine interfacing

Among the recorded units of each mouse/session, a set of 3 putative pyramidal neurons – the Master neurons – were selected by the operator. In the first two mice, we initially enrolled 7 neurons. However, after a first round of experiments, we found that securing so many large and high-firing neurons was challenging in several of the mice, so we settled on a smaller count of 3 neurons. We did not find any major difference in the activity or behavior of these first two mice. We selected the Master neurons among all simultaneously recorded units because they displayed (1) a sufficient baseline frequency (target: 10 Hz), (2) spikes clearly separate from the multiunit baseline and with the largest possible amplitude, and (3) a spike shape that was visually different from any other spike shape across the 4 channels of the tetrode.

The activity of these Master neurons was transformed into a virtual cursor position (Fig. 1C) which determined the optogenetic frame to be displayed as well as possible reward delivery. The spiking activity of the Master neurons was summed, and the corresponding firing rate was measured over 10 ms time bins. To transform this Master firing rate into the position of the virtual cursor, it was convolved with a 100 ms box kernel, and then renormalized with respect to the distribution of Master activity observed during a baseline window of 3 minutes just preceding the start of the session. Specifically, we computed the 99th percentile of the baseline activity values, and the activity from 0 Hz up to this value was split in 7 equal positions, with an additional 8th position for activity values exceeding the 99^th^ percentile threshold (Fig. S3). The resulting movements of the virtual cursor were smooth. In the Bar feedback condition, on average 95% of the transitions were to a closest neighbor position, and less than 0.1% of the transitions were jumps larger than to a second neighbor position. This was similar in all other feedback conditions.

For most of the experiments, only the 5^th^ position was rewarded, which means that whenever the virtual cursor was inside that position and the mouse simultaneously licked, it obtained a 5 μL water drop. Note that rewards were not delivered automatically to the mouse whenever the virtual cursor entered the rewardable position. Instead, only if the mouse licked at the precise time when the virtual cursor was located in the rewardable position, the capacitive sensor detected the lick and triggered the delivery of a drop of water through the lick port, which was immediately swept away by the ongoing licking action. In very first 3 experiments, only the 6^th^ position was rewarded, and in 3 additional experiments, the rewardable position also included either the 6^th^ or the 4^th^ position. We did not find any difference in activity or behavior that could be related to this difference in rewardable positions.

The logic of introducing a virtual cursor has been double. First, from a purely analytical point of view, it allows analysis of motor control in the non-linear discretized scale that is relevant for feedback stimulation and reward obtention, that is, regardless of the absolute values of firing rates which can be very different from one mouse and session to the next. Second, it emphasizes that the algorithm is the same in all feedback protocols. Only the final mapping between the eight different positions of the cursor and the effective photostimulation patterns changes with the protocol. This concept of a virtual cursor, in between the firing rate of the neurons and the photostimulation frames, is useful to describe unambiguously the protocols, the analyses and the results.

### Optogenetic photostimulation of somatosensory cortex

Each virtual cursor position was associated with a specific feedback pattern that was projected onto the barrel cortex of the mice using a Digital Light Processing module (DLP, Vialux V-7001, Germany). The DLP contained a 1024 x 768 pixels Texas Instruments micro-mirror chip, which was illuminated by a high-power 462 nm blue LED. The frame stream generated by the device was focused onto the cortical optical window using a tandem-lens macroscope (*58*), and covered the entire barrel cortex. We displayed each frame for 5 ms, followed by 5 ms without photostimulation. We sent homogeneous light spots, 225 μm in diameter, centered onto the barrel locations (see below). In a previous publication, we recorded activity in S1 in response to the exact same photostimulations, in the same mouse line, and verified that it triggered neuronal activation mostly limited to the targeted barrel area (Abbasi et al., 2018). In the same study we also compared the detection of five aligned spots flashed on the barrel cortex to the detection of five aligned spots flashed just outside the cranial window, in a GO/NOGO task. We found that mice detected the photostimulation only when it was targeted to the cortical window. This control ensured that the mice are unable to use any indirect clue, such as light reflection in the setup, to solve the task.

A set of at least 3 reference barrels, was localized on the mouse cortical surface via intrinsic signal imaging. These barrels were used to align a standard barrel map (*59*) that served later as the grid to align the photostimulation spots. Fig. S8 shows an example of the intrinsic signals and of the strategy used to position the photostimulations onto the S1 surface.

We used five different sets of feedback frames: the Bar feedback (Fig. 1D,E); three shuffled versions of the Bar feedback that are described in Fig. 3 and Fig. S4; and finally, a condition where no photostimulation was displayed (No feedback, all black frames). The Bar feedback design was based on the known selectivity of S1 neurons to features such as the global direction of bar-like stimulations (*26*, *27*, *60*) and more broadly, tuning to progressive movement of objects across the whiskerpad (*28*). This choice of feedback structure was also supported by the observation, in awake behaving rodents, that structured sweeping sequences of rostrocaudal deflections of whiskers are significantly more prevalent than expected by chance (*29*). Note that all photostimulation frames used the same number of identically shaped photostimulation spots, and therefore generated the same amount of photoactivation (*15*). The total amount of light projected onto the cortex was thus constant throughout all sessions.

To verify that the selected photostimulation did not bias the M1 activity prior to training, we exposed 3 naive, BMI-ready mice, to one single session of Bar feedback playback, and one of the Full shuffle playback. The frame sequence originated from a previous mouse/training session. During playback, in each mouse we recorded three M1 neurons that would qualify as Master neurons. We found no firing rate modulation triggered by any of the displayed frames (data not shown), and in particular none in the Bar feedback. These experiments confirm that prior to training, M1 neurons had no discriminative power or specific tuning to the photostimulation frames we designed.

### Behavioral training

We started the behavioral training by removing free access to the water in the cage. At the same time, we started habituating the mice to head fixation. This lasted for 2 days, where the mice were head-fixed during 30 min sessions and were continuously presented with a spout that delivered a drop of water every time the mice licked, thanks to a capacitive sensor in the spout.

After these first habituation sessions, we transitioned to training the mice in the BMI task. The sessions took place once a day, and lasted 33 min (including the 3-min baseline period). During these training sessions, the neuronal activity was continuously recorded, and one of the five photostimulation dynamical patterns was continuously applied to the mouse barrel cortex: Bar feedback, Barrel shuffle, Frame shuffle, Full shuffle, or No feedback, (Fig. 3A and Fig. S4). The displayed frame was updated every 10 ms based on the measured neuronal activity (Fig. 1). At any time, the mice could move the virtual cursor to the rewardable zone by modulating the activity of Master neurons. If it licked at the precise time when the virtual cursor was located in the rewardable position, a small amount (~5 μL) of water flowed immediately through the lick port, and the water droplet was swept away by the ongoing licking action.

We monitored the weight loss that resulted from the water restriction schedule. We ensured that through the whole training, the weight did not drop below 80% of its initial value, a consensus weight threshold in this model (*56*). To do so, mice were checked daily for weight and extra water/food intake was provided as needed to stabilize the weight. After these first sessions, we transitioned immediately to training the mice in the BMI task (1 session/day, 30 minutes), with one of the photostimulation feedback protocol, and only one feedback position rewarded.

The mice were trained with the same feedback protocol during 5 consecutive training sessions. There were no days off during these 5 days, except in the rare case of an unexpected technical problem. After the 5 training sessions, and if sufficient M1 activity was still present, we performed a new selection of Master neurons from scratch, and we restarted training the mouse with another feedback condition. There was generally a two days gap between different feedback protocols, except in three mice for which there was no pause in the training. We checked that previous learning did not bias the outcome of the following training (Fig. S5B).

If the recording of one neuron was lost during the training, the active Neighbor neuron with the largest spike shape was enrolled to replace it. If no additional Neighbor neuron was available, the experiment kept going with a reduced count of Master neurons, down to a minimum of two Master neurons. We assessed the Master neuron population stability by counting cases where all Master units could not be reliably identified anymore at the start of one of the training sessions, and had to be replaced with new units. This situation occurred once for two mice for the Bar feedback condition, two mice for the Full shuffle condition and three mice for the No feedback condition. This amounts to 7 cases out of 152 transitions between sessions, thus about 5%. Note that in most of these cases, although we were unable to prove it, we suspected that a least one of the former Master neuron was picked as part of the new Master neuron pool.

Videography of the snout in three mice failed to reveal whisker movements that would be correlated to optogenetic stimulation or to the virtual cursor motor control, as previously reported in BMI studies (*61*).

Through training, we monitored the weight loss that resulted from the water restriction schedule. We ensured that the weight never dropped below 80% of its initial value (*56*). To do so, mice were checked daily for weight and extra water/food intake was provided as needed to stabilize the weight.

### Offline spike sorting

Offline extraction of neuronal activity was performed using SpyKING CIRCUS (*62*). We confirmed that each online-sorted Master unit was properly spike sorted by matching it with a specific offline-sorted unit through comparison of spike shapes and amplitudes across tetrodes. All additional, non-Master offline-sorted units were labeled as Neighbor units.

### Data Analysis

All statistical tests are non-parametric, either two-sided Mann-Whitney tests or Wilcoxon paired tests.

### Intracortical microstimulation (ICMS) experiments

To confirm that the electrodes were located in the motor cortical area, we performed ICMS at the end of the behavior sessions (n = 3 mice, Fig. S2). We injected bipolar current pulses (amplitude 21 μA/channel, duration 1.4 s, frequency 60 Hz, 50% duty cycle) through the 32-channel NeuroNexus silicon probe implanted in M1, in awake head-fixed animals. The contralateral whiskers were imaged using a high-speed videography (camera – Baumer HXc-20, lens – 6 mm, F/1.4) at 300 frames per seconds for a duration of 9 s. A single trial consisted of 5 s pre-ICMS videography, followed by 1.4 s during ICMS stimulation and finally 2.6 s post-ICMS. This procedure was repeated 14 times during a single session of ICMS experiment, with a 1 s inter-trial delay. In the ICMS videos, a central whisker was identified amongst all the whiskers in the field of view, and tracked using the automated video tracking software DeepLabCut (*63*). The amplitude of ICMS-evoked whisker movement was defined as the mean whisker angle during the first 1 s of stimulation versus the 1s immediately before. Latency of whisker movement was measured at the first frame with significant whisker movement amplitude (2 standard deviations above the mean).

### Histology

After the experiment, mice were deeply anaesthetized with isoflurane (4-5%) and pentobarbital (150 mg/kg), then exsanguinated and perfused with 4 % paraformaldehyde (PFA). The brains were extracted and stored overnight in 4% PFA. The brains were then transferred to a solution of phosphate-buffered saline for at least 24 hours. Fifty μm slices were cut in the coronal plane and stained for cytochrome C oxidase. The location and depth of the silicon probe in the brain were traced by DiI depositing on the electrodes prior to their implantation and by localizing afterwards the fluorescent dye present in the histological slices (Fig. S2A).

## Acknowledgments

We thank Isabelle Férézou for advice on experimental design and analysis; Ehud Ahissar, Boris Barbour, Karim Benchenane, Suliann Ben Hamed, Sliman Bensmaia, Daniel Feldman, Evan Harrell and German Sumbre for comments on an earlier version of the manuscript. Aurélie Daret and Guillaume Hucher provided experimental support.

## Author contributions

DES and VES obtained funding and administrated the project. DES, LE and VES conceptualized and supervised the experiments and analysis. AA and HL carried the investigation with support from LE and DG. LE and AA analyzed the data with support from HL. LE, VES and DES wrote the manuscript with input from all authors.

## Data and materials availability

All data and code are available upon reasonable request to the corresponding author.

