## Supplementary Figures for "Brain-machine interface learning is facilitated by specific patterning of distributed cortical feedback"

**This PDF file includes:**

Figs. S1 to S8

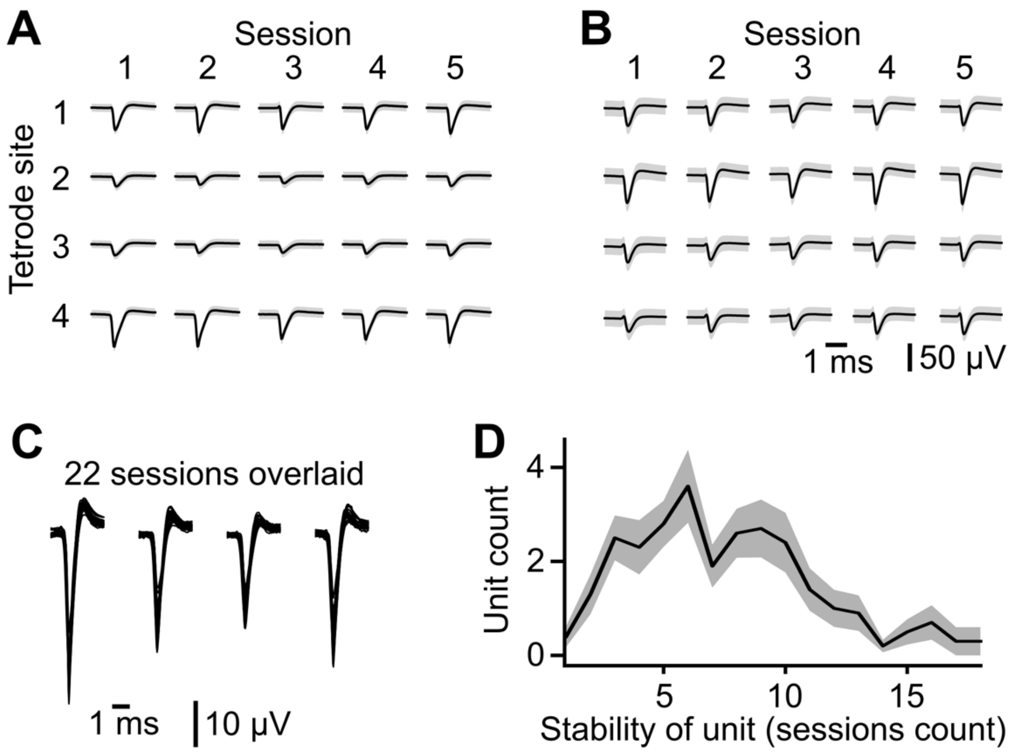


**Fig. S1. Stability of the recorded neurons across sessions. (A)** Stability of the spike shape of one Master neuron, shown for the 4 tetrode sites, across five training sessions, in the No feedback condition. **(B)** Same as a for a second Master neuron, this time in the Bar feedback condition. **(C)** Overlay of the spike shapes for one Master neuron across 22 successive sessions. **(D)** Average distribution across 10 mice of the count of sessions where the same Master unit could be identified. Shaded background: standard error of the mean of the histogram.

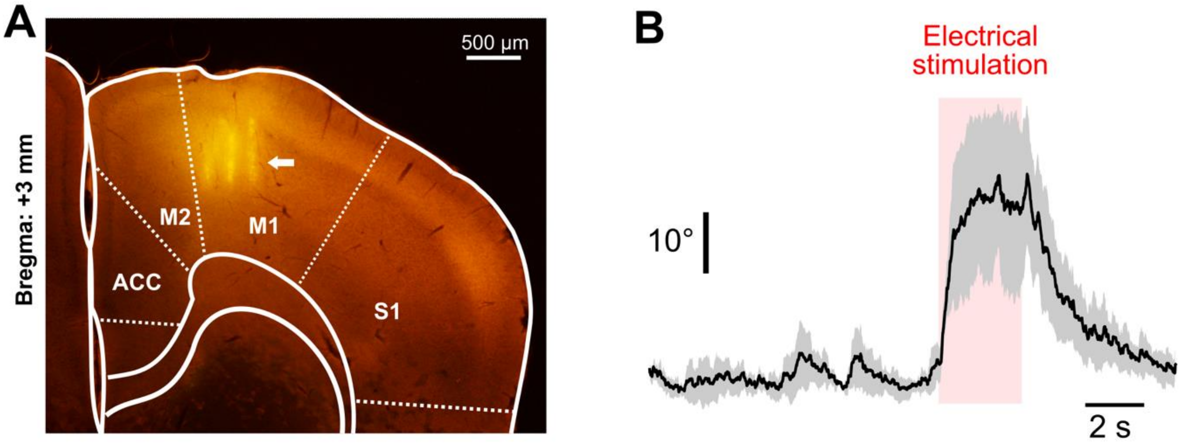


**Fig. S2. Localization of the implanted silicon probes in whisker M1**. **(A)** 50 µm coronal slice of a mouse brain, stained for Cytochrome oxidase. DiI coating of the shanks prior to insertion resulted in fluorescent lines indicative of the location of single shanks (yellow tracks) in M1 (white arrow). Dashed lines: area borders according to the Allen brain atlas. We identified the electrodes as being placed in the deeper layers of M1 based on: the location of the slice with respect to bregma; the lateral location of the electrode tracks with respect to the longitudinal fissure; and the depth of these tracks. **(B)** The amplitude of angular movements of a contralateral whisker evoked by ICMS stimulation through the silicon probe (60Hz, 21 microA, average of 3 mice) confirms that the electrode was located in M1 (see Methods). Shaded background: standard error of the mean.

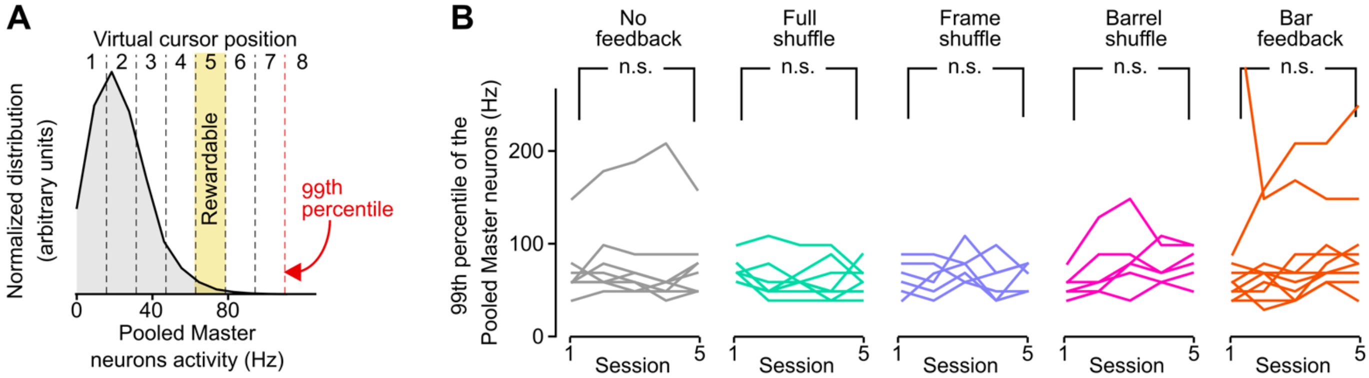


**Fig. S3. Firing rate limits defining the eight possible positions of the virtual cursor. (A)** Example distribution of the pooled firing rate of Master neurons during the 3 min baseline at the start of one session. The transition between virtual cursor positions 7 and 8 is set at the 99th percentile of the firing rate distribution. The range from 0 Hz to the 99$^{th}$ percentile is divided into equal firing rate intervals. Each firing rate interval corresponds to one virtual cursor position from 1 to 7 as indicated. **(B)** The 99th percentile of the pooled Master neurons firing rate did not evolve significantly over training sessions, regardless of the protocol. Each line: 1 mouse. (n.s.: Mann-Whitney p value > 0.05).

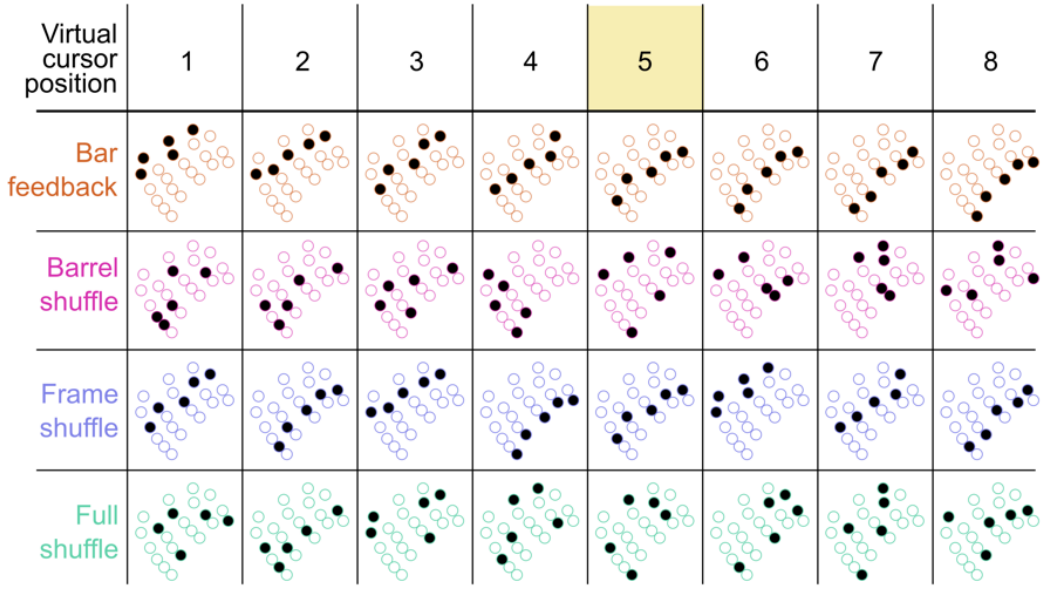


**Fig. S4. Detail of the photostimulated barrels in the four feedback conditions, across the eight virtual cursor positions.**

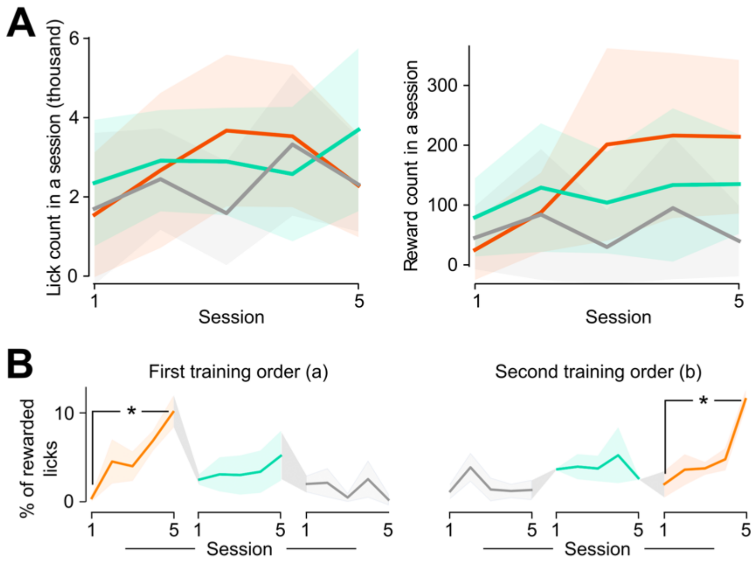


**Fig. S5. Engagement and performance during training across feedback conditions. (A)** Engagement of the mice during training sessions in the Bar feedback (orange), Full shuffle (green) and No feedback (gray) conditions. Left: total number of licks per session. Right: total number of rewards per session. Lines: average across mice. Shaded backgrounds: ± SEM. **(B)** Learning curves for two groups of mice (n = 3 in each group) trained with two different orders of presentation of three protocols: Bar feedback (orange), Full shuffle (green) and No feedback (gray). *: $p < 0.05$. Mann-Whitney non-parametric tests. Lines: average across mice. Shaded backgrounds: ± SEM.

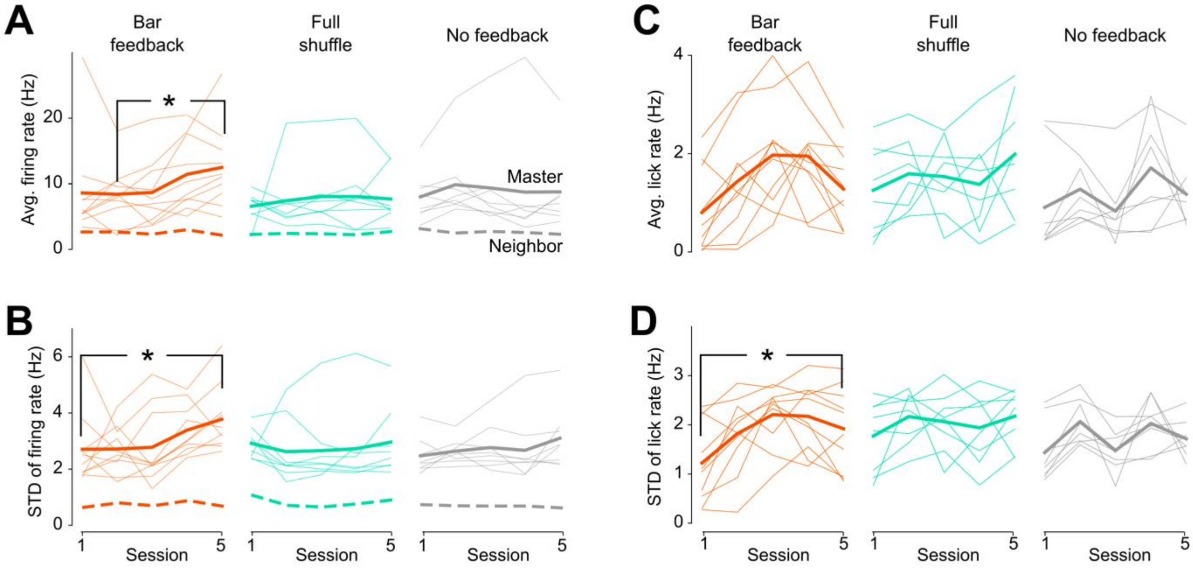


**Fig. S6. Firing statistics of Master and Neighbor neurons, and licking statistics, during closed-loop learning in 3 feedback conditions. (A)** Thin lines: Firing rate averaged over the individual Master neurons of one mouse as a function of training sessions. Thick lines: Average of individual neuron firing rates across all mice, for Master (continuous line) and Neighbor (dashed line) neurons. Bar feedback (orange), Ful shuffle (green) and No feedback (black) conditions. **(B)** Same as A, for the standard deviation of the firing rate (measured over 1 s windows). **(C)** Average lick rate over sessions. **(D)** Average standard deviation of the lick rate (measured over 1 s windows). All tests: Mann-Whitney. *: $p < 0.05$.

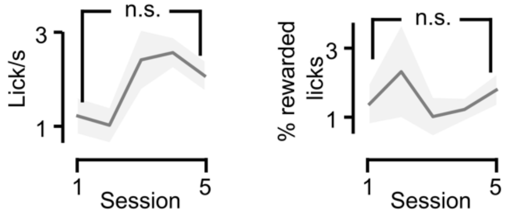


**Fig. S7. Absence of learning during 5 sessions of bar playback training in five mice.** Left: Mean (+/-SEM) frequency of licking per session. The increase was not significant. Right: Average (+/-SEM) proportion of rewarded licking over the 5 training sessions. The modulation is not significant. n.s: Wilcoxon test p>0.05.

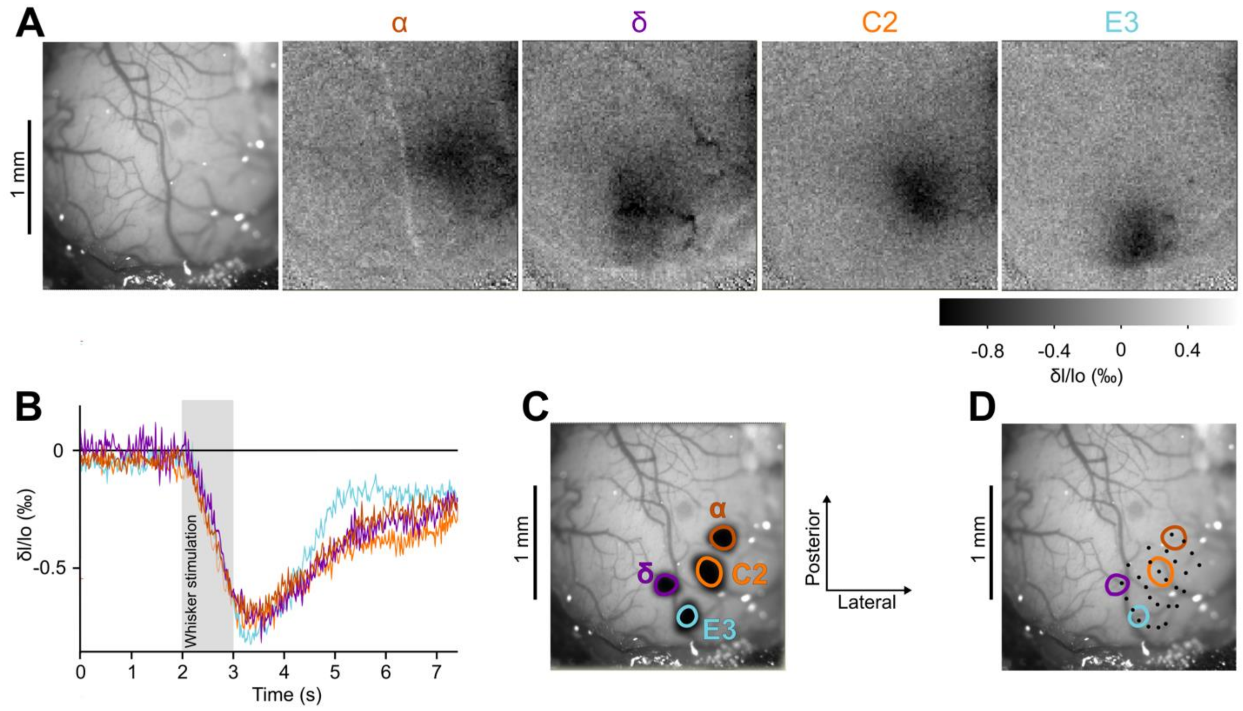


**Fig. S8. Identification of the barrels position within the chronic optical window using intrinsic imaging shown for one mouse. (A)** Image of the barrel cortex showing the blood vessels and intrinsic imaging during stimulations of the Alpha, Delta, C2 and E3 whiskers. **(B)** Time course of the whisker stimulation and of the intrinsic signal. Each whisker was stimulated for 1 second (grey area) with 100 Hz rostro-caudal deflections. **(C)** Thresholded contours of the intrinsic signal peaks used to define the location of the barrels. Contours of barrels correspond to 85% of the maximum relative absorption, after applying a 20$^{th}$ order gaussian filter. **(D)** Alignment of the barrel map from Knutsen et al., 2016, with the 4 barrels localized by intrinsic imaging.
